# Deep learning classification of *ex vivo* human colon tissues using spectroscopic OCT

**DOI:** 10.1101/2023.09.04.555974

**Authors:** Wesley Y. Kendall, Qinyi Tian, Shi Zhao, Seyedbabak Mirminachi, Abel Joseph, Darin Dufault, Chanjuan Shi, Jatin Roper, Adam Wax

## Abstract

Screening programs for colorectal cancer (CRC) have had a profound impact on the morbidity and mortality of this disease by detecting and removing early cancers and precancerous adenomas with colonoscopy. However, CRC continues to be the third leading cause of cancer-related mortality in both men and woman, partly because of limitations in colonoscopy-based screening. Thus, novel strategies to improve the efficiency and effectiveness of screening colonoscopy are urgently needed. Here, we propose to address this need using an optical biopsy technique based on spectroscopic optical coherence tomography (OCT). The depth resolved images obtained with OCT are analyzed as a function of wavelength to measure optical tissue properties. The optical properties can be used as input to machine learning algorithms as a means to classify adenomatous tissue in the colon. In this study, biopsied tissue samples from the colonic epithelium are analyzed *ex vivo* using spectroscopic OCT and tissue classifications are generated using a novel deep learning architecture, informed by machine learning methods including LSTM and KNN. The overall classification accuracy obtained was 88.9%, 76.0% and 97.9% in discriminating tissue type for these methods. Further, we apply an approach using false coloring of *en face* OCT images based on SOCT parameters and deep learning predictions to enable visual identification of tissue type. This study advances the spectroscopic OCT towards clinical utility for analyzing colonic epithelium for signs of adenoma.

## 1 INTRODUCTION

Colorectal cancer (CRC) is the third most diagnosed cancer in the United States among both men and women, with approximately 153,020 new cases expected in 2023 [1] It is also the second leading cause of cancer deaths in the U.S. with over 52,000 expected to die from CRC each year [1]. Early detection of CRC as well as detection and removal of precancerous polyps (i.e., adenomas) via screening colonoscopy and other modalities has led to reduced mortality rates [2, 3]; yet, less than half of adults undergo regular screening [4] and the rate of cancer incidence has plateaued in the past decade among adults ages 50-64 [5]. More concerning is a steady increase in incidence of those aged 49 years or less [5]. The recent decline in effectiveness of CRC screening is due to several factors, including the use of subjective biomarkers and lack of real time feedback to the endoscopist. Polyps found during colonoscopy are resected and evaluated via histology to identify malignancies. Even though colonoscopy is an effective technique, there is room to improve efficiency. Recent efforts in endoscopic innovations seek to improve efficiency by reducing procedure time, with its associated risks, and pathology costs by leaving diminutive polyps (<5mm) in place. These small lesions account for 80% of resected polyps during colonoscopies [6]. If a leave-in-situ strategy was successful, it could realize millions in cost savings by reducing procedure time while also reducing the subsequent cost of processing and analysis [7].

OCT has been explored as a means for optical biopsy since early in its development [8] for its ability to obtain cross sectional images of tissue noninvasively while also providing label free microscopic resolution. However, the technique only provides structural information without the functional information needed to evaluate tissues for early signs of cancer. One avenue for augmenting OCT is to use spectroscopic information. In spectroscopic OCT (SOCT), a short time Fourier transform (STFT), i.e. a windowed transform, is used to analyze the wavelength dependence of OCT signals [9, 10] and reveal optical tissue properties.

SOCT was first applied to colonic epithelium in a study to detect cancer development in the azoxymethane (AOM) rat carcinogenesis model [11]. In this study, SOCT was used to analyze tissue optical properties and determine the size of cell nuclei as a function of depth into the tissue. Another approach, termed inverse spectroscopic OCT (ISOCT) uses a rigorous model of tissue scattering to determine optical properties from SOCT data [12, 13]. The ISOCT approach was used to discriminate advanced adenoma in colorectal epithelium from biopsy samples. Most recently, the Zhu group has found that colorectal epithelium could be evaluated using maps of scattering coefficient [14, 15], an optical property measured with OCT. Extending their approach using deep learning also showed improved diagnostic capacity [16].

In this work, SOCT is used to evaluate biopsies of colonic epithelium to quantitatively assess optical properties across a range of histology. Two optical properties of scattering are used in tandem, attenuation coefficient and scattering power, to evaluate the potential for monitoring CRC development. The optical property data is used to develop machine learning and deep learning models to distinguish tissue types. We have previously used this approach to evaluate CRC in a mouse model [17]. Here we extend this analysis to examining human colonic epithelium. We show that optical properties can be used as a source of information for false coloring schemes which can be used to visually identify tissue structures. The optical property data are then used to train machine learning models which show potential utility for automated analysis of tissue health.

## 2 METHODS

### 2.1 Instrumentation

The OCT system used in this study was based on a modified version of the OQ Labscope (Lumedica, Inc., Durham, NC) shown in Figure 1. Two superluminescent diodes (Superlum, Ireland) were used to provide a broad bandwidth. The combined source provided a center wavelength of λ_o_ = 903 nm, with a FWHM bandwidth of 180 nm, producing a axial resolution of 1.4 microns. The spectrometer contains a 2048-pixel CCD line array camera (OctoPlus, Teledyne e2v), with a line rate of up to 80,000 per second. The lateral resolution was set at 5 microns by using a 10X microscope objective attached to the end of the scanner. The motorized reference arm uses a pair of mirrors on a translating stage to vary the optical pathlength and an electronically controlled liquid lens from Optotune (Dietikon, Switzerland) to control the amount of light that is coupled back into the interferometer. The stage provides a physical pathlength range of 30 mm which gives 60 mm of optical pathlength change since it is a double pass.

**Figure 1:**
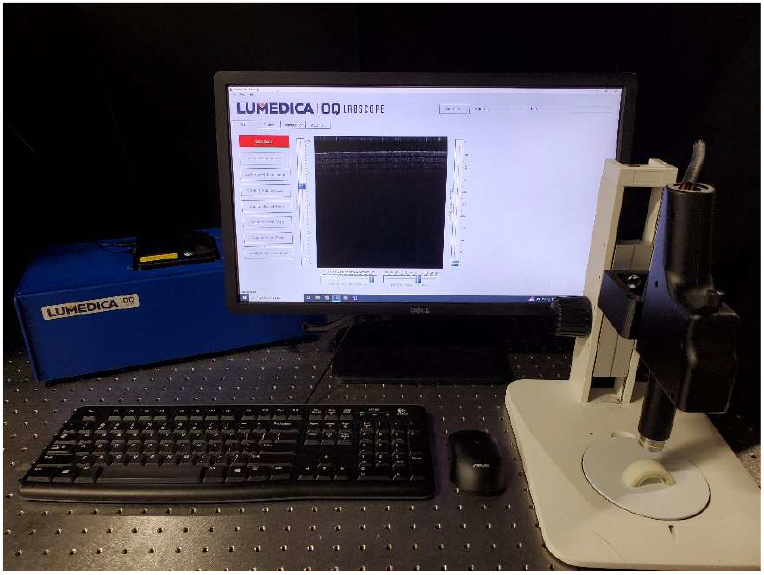
Photo of Lumedica (Durham, NC) OQ Labscope system used for this work. OCT engine (blue box) contains optical components and integrated PC. Scan head is mounted on a microscope stand for imaging samples.

### 2.2 Spectroscopic OCT and optical properties

Spectroscopic optical coherence tomography (SOCT) is a functional extension of OCT that obtains spectral information using a windowed Fourier transform, also known as the short time Fourier transform (STFT) in analogy to time domain signals [9]. Spectral domain OCT is based on using a broadband light source in an interferometry scheme and spectrally resolved detection, enabling direct post-processing means to extract the spectral properties of the sample. Instead of simply using a Fourier transform to translate the measured spectral data into a depth resolved reflection profile, the windowed Fourier transform produces such profiles as a function of wavelength [13]. The use of the STFT causes a reduction in axial resolution. Although the dual window method has shown the ability to avoid this trade-off [18], it is not used here where axial resolution is not a priority. From the spectral data, two optical properties are extracted from the SOCT data to classify tissues, the scattering attenuation coefficient μ_s_ as well as the scattering power (SP). A visualization of how these coefficients are obtained is shown in Figure 2

**Figure 2:**
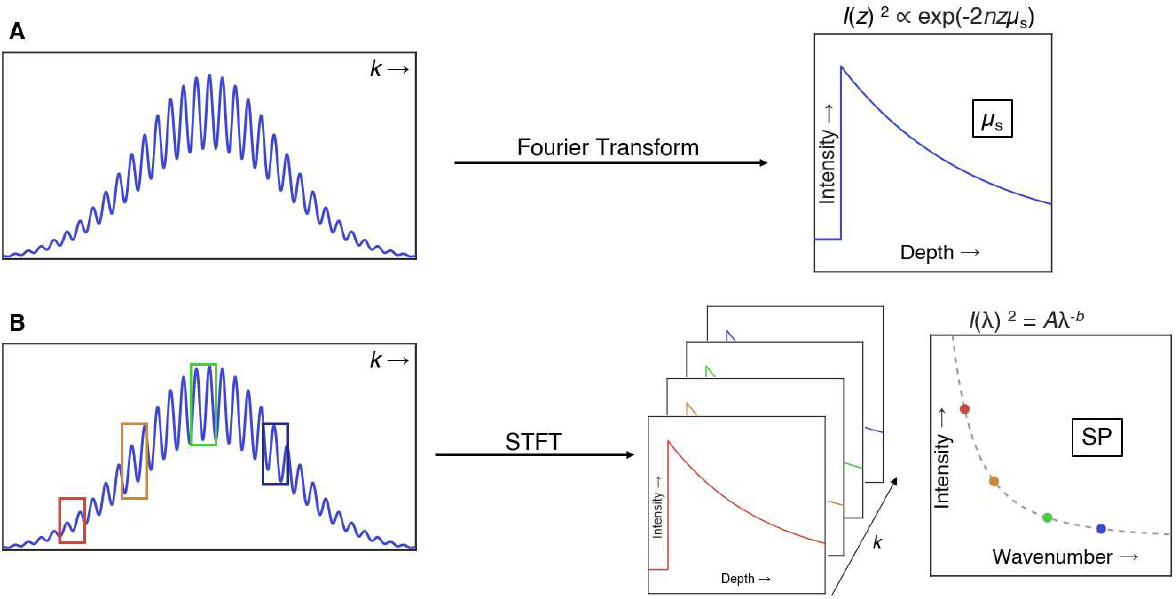
Visual representation of data processing steps performed to obtain optical coefficients. (A) Fourier transform of spectral domain OCT data produces a depth resolved reflection profile (A-scan) which provides the scattering attenuation coefficient μ_s_. (B) Windowed Fourier transform applied to produce SOCT data. Each window, represented by colored boxes, can be used to generate an A-scan for the selected frequency. The spectral variation across these scans is fit to a power law exponential to obtain the scattering exponent *b* which is used to determine the scattering power (SP = 4 – *b*). Adapted from [17]

Each OCT depth resolved reflection profile (A-scan) typically exhibits an exponential decay, as shown in Fig. 2 (a). The magnitude of the A-scan depends on the backscattering coefficient while the dependence with depth is characterized by the scattering attenuation coefficient μ_s._ [19]. In general, the scattering attenuation coefficient varies with wavelength. Using SOCT, this dependence can be measured and analyzed, as shown in Fig. 2 (b). The spectral dependence is modeled using a power law [20] Aλ^-b^, where A and b are fitted parameters, and 4 - b is the scattering power (SP) [21-23].

### 3.2 Human Clinical Study

A human clinical study was performed at the Duke Endoscopy Clinic with approval from the Duke University School of Medicine IRB. The study was designed to obtain the optical properties from a wide variety of adenomas in the colon. By including a diverse set of adenomas with different pathologies, an atlas can be created to obtain optical coefficients for specific adenoma types as input for developing classification algorithms. OCT imaging data were collected from 19 patients, consisting of volumetric scans of freshly excised adenomas. The protocol was modified after patient 11 to include normal mucosa tissue samples from the colonic epithelium for comparison.

Inclusion criteria included those who are 18 years or older, able to provide consent, and were referred for colonoscopy at the Duke University Hospital Division of Gastroenterology for resection of previously identified colon polyps or multiple polyps. Exclusion criteria included a history of coagulopathy, high international normalized ratio (INR) > 2, and low platelet count (less than 50 x 10^9^/L), or lack of “lift sign” indicating the presence of a cancerous lesion that must be removed via alternative procedures. Patients meeting the inclusion criteria and enrolled in the study underwent colonoscopy where their polyps were resected using endoscopic mucosal resection (EMR) or endoscopic submucosal dissection (ESD). After the polyps were removed, they were immediately imaged *ex vivo* using the SOCT instrument, followed by fixation in formalin for standard of care histopathological analysis. Additionally, 6 biospies of normal-appearing colonic eptihelium, located approximately 20 cm from resected polyps are also removed using forceps., These normal tissue samples were also immediately imaged using the SOCT instrumentation.

## 4 RESULTS

### 4.1 Human colon biopsy data set

A summary of the patient data broken down by tissue morphology is given in Table 1. Imaging data from 8 patients were selected for analysis to include the same number of tubular and tubulovillous samples (4 each). Normal mucosa tissue data for two of these patients was not available as described above. The data were separated into three groups: 90% training, 5% validation, and 5% testing using randomly selected tissue locations within each volume scan. The total number of A-scans used in this study were: Tubular: 783,353, Tubulovillous: 757,468, Normal mucosa: 764,481, Total 2,307,302.

**Table 1:** Overview of patient information from clinical data, separated into training and test data. Check indicates data that were acquired, x-mark indicates data were not available.

| # | Benign Tissue | Polyp | Morphology |
| --- | --- | --- | --- |
| 8 | X | ✓ | Tubulovillous |
| 9 | X | ✓ | Tubulovillous |
| 12 | ✓ | ✓ | Tubulovillous |
| 16 | ✓ | ✓ | Tubulovillous |
| 11 | ✓ | ✓ | Tubular |
| 14 | ✓ | ✓ | Tubular |
| 17 | ✓ | ✓ | Tubular |
| 19 | ✓ | ✓ | Tubular |

A variety of information was collected about each patient in case it would be potentially useful for later classification. Information included presence of dysplasia, mechanism of polyp removal, and polyp bulk and microstructural morphology. Normal mucosa tissue imaging data were limited because some patients consented to imaging of polyps but specifically declined consent for imaging of normal mucosa tissue bites. Because of the wide diversity of patients enrolled, it was decided that emphasis would be put on using data from patients with both normal mucosa tissue and polyps in training the algorithms to eliminate any factors unique to the patients themselves from influencing classification.

Classifications of “normal mucosa”, “tubular”, and “tubulovillous” are used as distinct morphological descriptors for the dataset, with each A-scan and the particular optical properties (spectrum sequence data and μ_s_ parameters) used in the analysis. Information about location within the colon where the polyp was found as well as the mechanism of polyp removal were not used as training data. Some patient data were excluded from the set to balance out the number of A-scans of each morphology available.

Data consist of volumetric data taken at multiple polyp locations from the patients, no distinction was made between different locations on the same polyp vs. different polyps as long as they came from the same patient. Certain volume scans were deliberately withheld from training to serve as test data with the aim of later determining the accuracy of the produced classifiers on previously unseen polyps from the same patients versus previously unseen polyps from new patients.

### 4.2 SOCT parameters for tissue contrast

To assess feasibility of using SOCT for visualizing tissue structure, μ_s_ and scattering power values were obtained from volumetric scans of *ex vivo* human colon polyps. While these two parameters were not sufficient in providing good classification of polyp type alone, it was found that μ_s_ reveals subsurface structural information about the tissues that is not visible in the topographical information. Figure 3 shows an example of this type of contrast. The left panel shows the tissue profile obtained using OCT intensity information. The general shape of the biopsy section is visible with darker blues signifying a greater tissue height. The middle panel shows the contrast gained from using μ_s_ as the contrast mechanism. In this figure, the glandular structures are visible with the greater yellow hue indicating higher μ_s_ values. The dark horizontal lines are due to signal dropout arising from hyperreflections.

**Figure 3:**
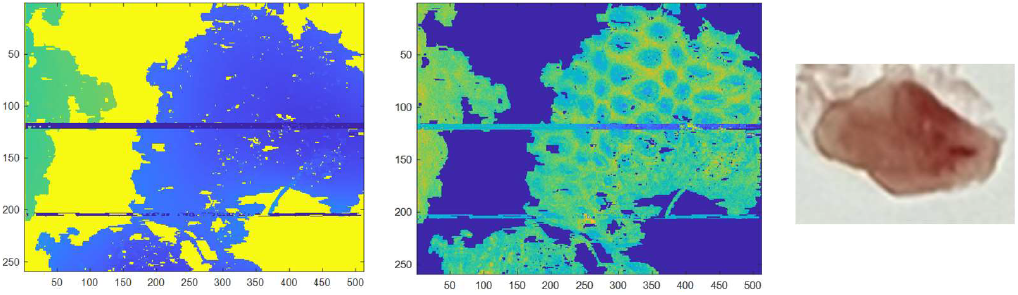
(Left) En face volumetric OCT scan of a normal mucosa tissue biopsy. False coloring is applied based on tissue topography, with taller heights corresponding to darker blue. (Middle) Same en face volumetric OCT scan but with a colormap indicative of the value of *μ*_s_ at individual biopsy locations. Higher *μ*_s_ values correspond to more yellow pixels. A crypt structure in the tissue can be clearly seen from the *μ*_s_ information that is not visible from the topographical edge map alone. (Right) Gross pathology of the imaged tissue biopsy.

Figure 4 shows how this optical property can be used to discriminate tissue types using a biopsy of an adenomatous tissue. Again, the OCT tissue map shows the outline of the tissue biopsy (left) while the map generated with μ_s_ values reveals a subsurface tissue architecture (middle). A more disordered crypt like structure is apparent in this tissue, elucidated by the information in μ_s_.

**Figure 5:**
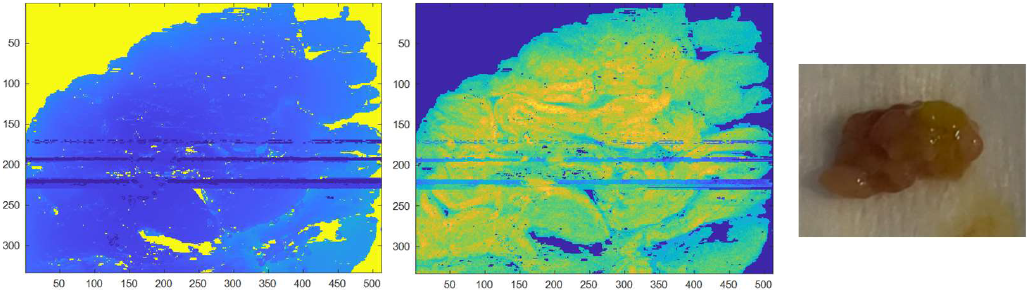
(Left) En face topographical map of the volumetric OCT scan for adenomatous polyp. (Middle) Map of polyp with a false color overlay based on *μ*_s_ values. (Right) Gross tissue pathology.

Interestingly, the quality of the fit used to generate the μ_s_ maps can also be used as a source of contrast. By plotting the r-squared value for the quality of each fit tissue structure is revealed. Figure 6 shows examples of this imaging using one normal mucosa tissue sample (left) and two polyps (middle, right). The corresponding H&E stained tissue micrographs are shown These images provide similar information to the μ_s_ false color overlays with even higher contrast. The heterogeneity of epithelial tissue may result in poorer quality linear fits due to subsurface layers attenuating light differently; higher r-squared values indicate a better fit, suggesting a more homogenous subsurface structure. Notably, these changes can be observed at high resolution: structural changes are observed without any binning of neighboring A-scans, resulting in a 15 micron lateral resolution for examining structural differences within the bulk tissue, which approaches cell-level resolution.

**Figure 6:**
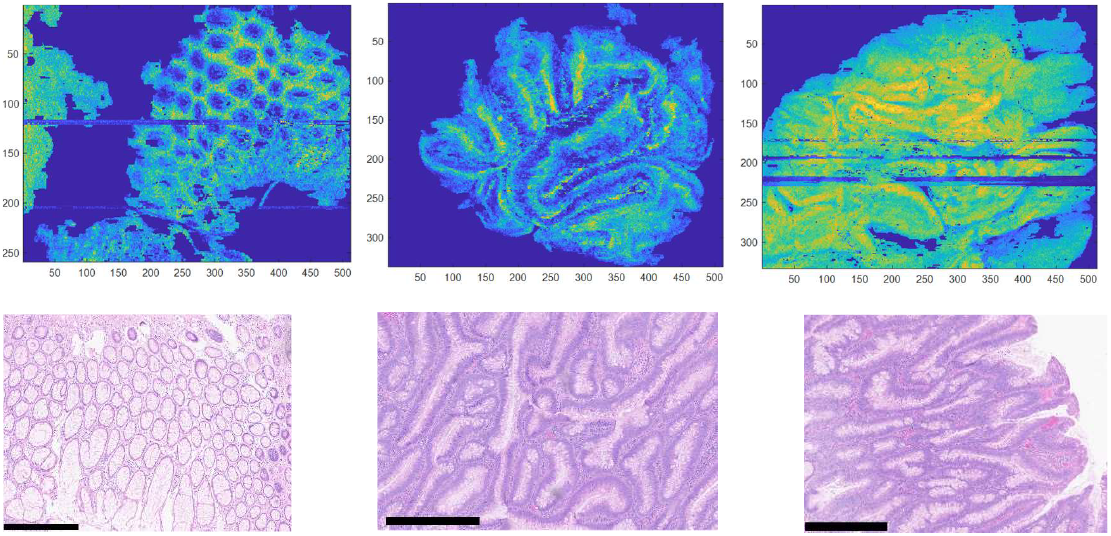
En face volumetric OCT scans with false color overlay representing r-squared values for fitting to mus curves. Yellow hue indicates higher r-squared value. These images look similar to those from the *μ*_s_ overlay but with increased contrast.(Left) Normal mucosa and (Middle, Right) adenomatous tissue biopsies with corresponding pathology beneath.

### 4.2 Preliminary analysis used for classification approach

In order to develop the deep learning classifier, preliminary analysis was conducted using machine learning methods. This approach enabled discovery of the spectroscopic information that could best discriminate tissue type. Based on previous work in the mouse colon [17], we used a bidirectional LSTM network to study the optical property data from four patients, from whom both polyps and normal mucosa tissue samples were collected. Approximately 15 percent of the A-scans were randomly chosen as test data and the remainder used for training data. Similarly to the mouse imaging study[17], μ_s_ alone did not produce meaningfully accurate discrimination but scattering power alone was useful and produced results with fairly high accuracy. The data set was comprised of 72,440 A-scans from the four polyp biopsies and 37,728 from the normal mucosa tissues. The use of scattering power alone resulted in an overall accuracy of 90.8% with a sensitivity of 89.2% and a specificity of 94.9% using the LSTM network. This result suggested that scattering power offers some utility but with room for improvement, particularly by adding μ_s_ information.

The bidirectional LSTM network is based on assuming that the optical property data has an underlying functional relationship between adjacent data points. Although this approach showed some success for the spectral data, the relationship between adjacent depth points was not strong enough for μ_s_ on its own to provide discriminatory capacity using this network. To investigate what aspects of μ_s_ information may be able to provide discrimination of tissue type, a basic KNN-based model was developed based on μ_s_ and parameters derived from it.

Each parameter was examined for a 3x3 grid of adjacent A-scans. The nine features included the mean value of μ_s_, the mean r-squared value associated with the linear fit, the mean RMSE value associated with the linear fit, the standard deviation of μ_s_, the correlation coefficient across the A-scans, mean peak intensity, mean of the mean intensity (across all nine A-scans), mean of the standard deviation of intensity (across all nine A-scans), as well as the maximum height. Any 3x3 grid point was omitted if any pixels contained an edge profile or signal drop out. After the KNN was trained, the overall validation accuracy was 76.0%, with 78.9% sensitivity and 72.3% specificity. Although these data do not show an improved accuracy compared to the spectral data and the LSTM approach, the features could be examined for relevance.

Figure 7 shows a bar graph representing the minimum redundancy maximum relevance (MRMR) feature selection output. Interestingly, the μ_s_ values themselves were found to be one of the least influential features. This provides insight into why efforts to develop discrimination algorithms based solely on these values were unsuccessful. The most influential features are seen to be those derived from the intensity (the peak A-line intensity, the mean intensity of the A-lines and the standard deviation of intensity of the A-lines) as well as the goodness of fit of the depth profile (r-squared and RMSE). The use of KNN for classification points the way to using a combination of features related to μ_s_ to discriminate between the different tissue types. However, a limitation of the KNN method is that it is not a deep learning method, meaning that it cannot be pre-trained for analyzing data. Instead, we use the lessons learned from the KNN-based analysis as a step towards developing a deep learning approach.

**Figure 7:**
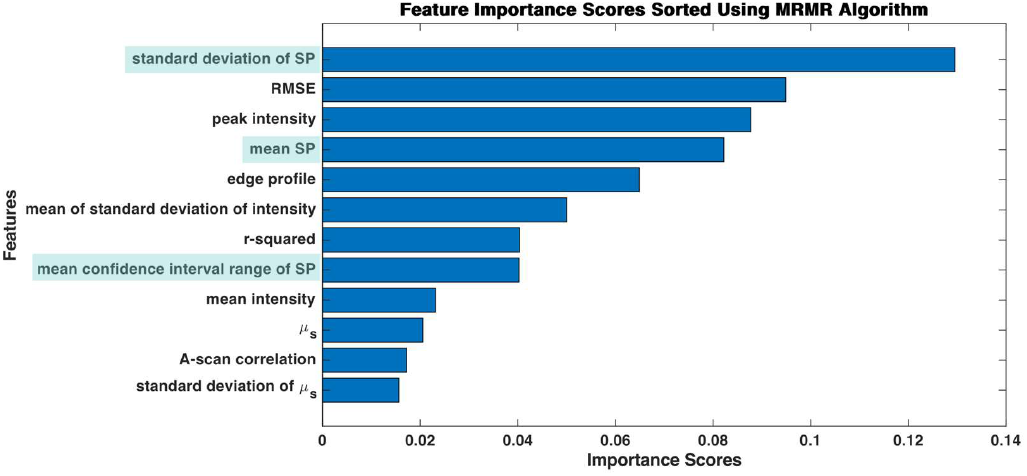
Bar graph showing the relative importance of specific *μ*_s_ and SP-related features as determined by a minimum redundancy maximum relevance (MRMR) feature selection algorithm from MATLAB.

### 4.4 Tissue classification using deep learning

A neural network classifier was developed to discriminate tissue type from the parameters derived from the SOCT data (Figure 8). As with the KNN approach, the data were binned into 3x3 grid of A-scans to limit the variability from scan to scan and provide an effective area for an optical biopsy. The nine μ_s_ features mentioned above were used as scalar valued features in the feature input layer along with three additional features from the spectrum: mean scattering power, standard deviation of scattering power, and mean 95% confidence interval for power law slope fitting. These data are passed through three fully connected layer blocks, each consisting of a fully connected layer, a batch normalization layer, and a leaky ReLU layer.

**Figure 8:**
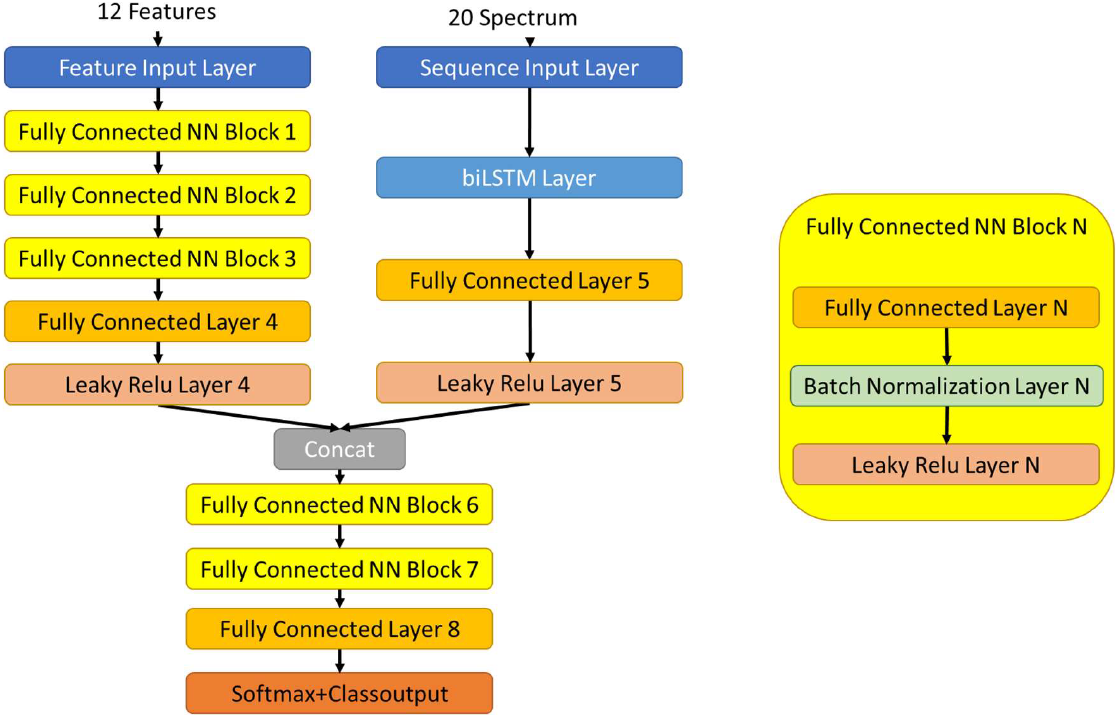
Schematic of the network architecture for the fully connected neural network developed for analyzing human colon tissue data. The scatter power data uses a similar biLSTM layer to that described above, while the μ_s_ data is trained on a fully connected structure, with the two outputs concatenated together.

The neural network is supplemented using additional spectral information derived from the bidirectional LSTM discussed above. A separate sequence input is created based on the data from the biLSTM layer which is then passed through fully connected and leaky ReLU layers. The information from these two pathways is combined using a concatenate layer that reshapes all of the inputs into a single tensor which is then passed through two more fully connected layer blocks before being input to a softmax activation function to give a single output decision of normal mucosa, tubulovillous, or tubular. This architecture combines the discrimination obtained by using the KNN to analyze tissue data based on scattering attenuation coefficient, as well as the spectral data that yielded discrimination via the power law using the biLSTM layer. The algorithm comprises a total of 26 layers and 104,300 learning parameters. The alpha-value for the leaky ReLU layers was 0.01, and 100 hidden layers are included within the biLSTM layer. The data were split such that 90% was used for training, 5% for validation and 5% for testing. The batch size was 128 and training was conducted over five epochs. The Adam optimizer was used with β_0_ = 0.9, β_1_ = 0.999, and ε = 10^-8. Training was implemented with an Intel i7-9750H processor, and total training took 115 minutes.

Figure 9 summarizes the classification performance of the neural network applied to the test data (the 5% of scans which were withheld from training) for distinguishing the tissue samples as either normal mucosa, tubular, or tubulovillous adenoma. Individual biopsy level classification accuracy was high. There was 96.04% accuracy in identifying normal mucosa tissue, with 0.35% of normal mucosa scans being misclassified as tubular adenomas and 2.27% misclassified as tubulovillous adenomas. There was 92.64% accuracy in identifying tubular adenomas, with 0.34% of tubular scans being misclassified as normal mucosa and 8.51% of tubular scans being classified as tubulovillous. Finally, there was 90.78% accuracy in identifying tubulovillous adenomas, with 3.27% being misclassified as normal mucosa and 5.79% misclassified as tubular. The overall classification accuracy was 93.15%. A histological-based dichotomy is also considered for sensitivity and specificity measurements. Under the histological dichotomy where normal mucosa tissue is separated from tubular and tubulovillous tissue, the sensitivity is 98.21% and the specificity 95.21%, with an overall accuracy of 97.92%.

**Figure 9:**
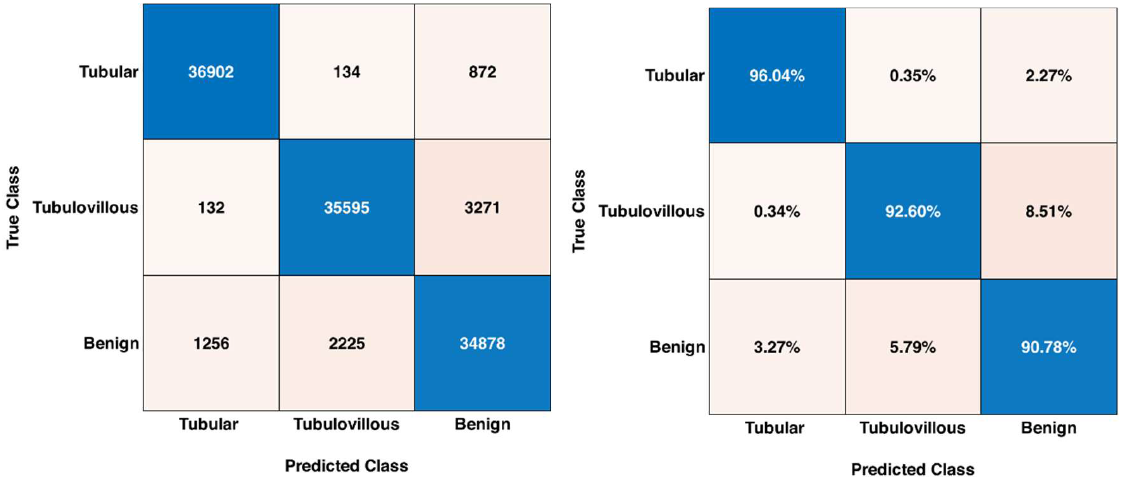
Confusion matrix showing results using the developed deep learning architecture. (Left) Individual biopsy level selections, (Right) Percentages broken down by prediction.

### 4.5 Visualization of tissue classifications

In order to demonstrate a potential approach for using the network output for informing clinical decisions, we now present its application to three volumetric OCT scans which were deliberately withheld from the training set. The classifications are now presented visually with probability output from each channel (normal mucosa, tubular, tubulovillous) first scaled to a value between 0 and 255 and then assigned to an RGB color channel as shown in Figure 10.

**Figure 10:**
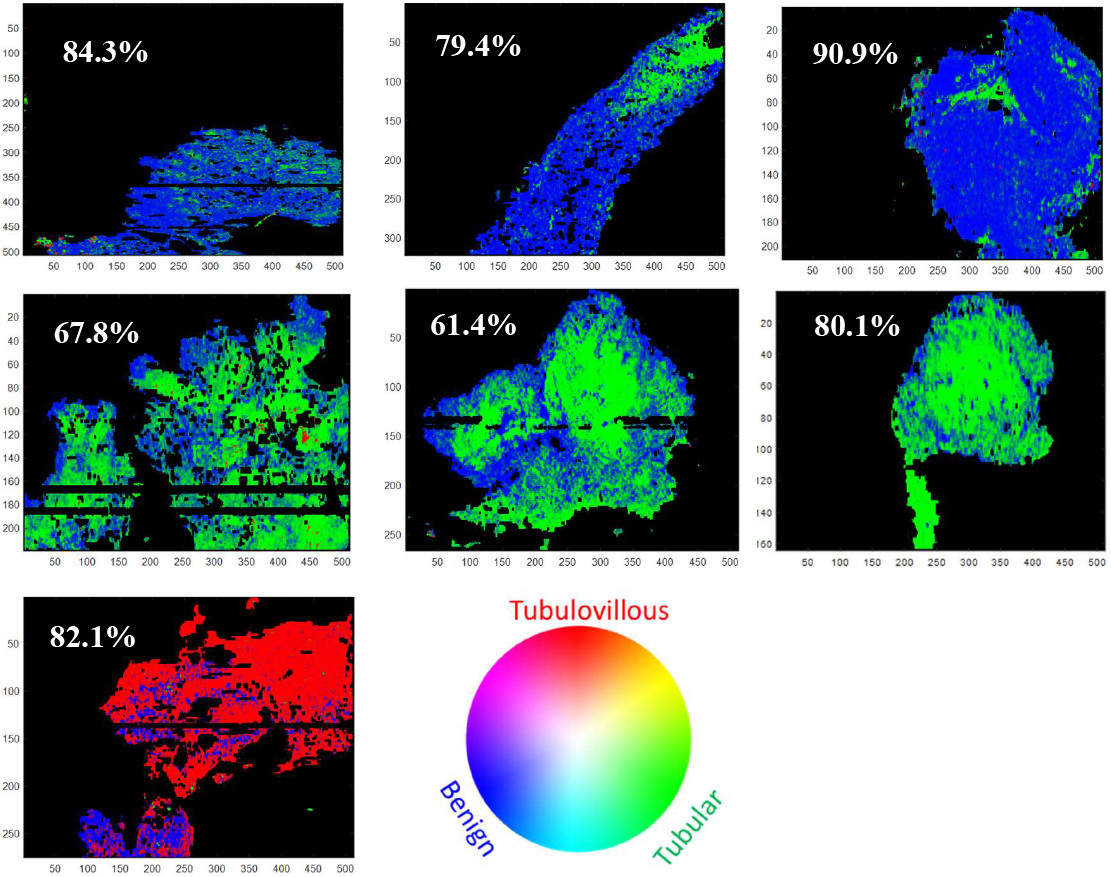
OCT images overlayed with false color indicating classification results per the color wheel. Top row shows images from normal mucosa (benign) tissue samples, middle row shows results from tubular tissue samples, the image in the bottom row is for tubulovillous tissue sample.

These data were from patients where imaging data was used in the training dataset but these specific scans were not used in the training. Overall, this approach had good accuracy in identifying each optical biopsy (3x3 binned A-scans) ranging from 61% to 91%.

Although the accuracy of each individual optical biopsy can be somewhat variable when compared to the overall accuracy seen in testing of the neural network, the distribution of predictions across a given tissue sample provides a good indication of the overall disease state. The incorrect classifications for individual optical biopsies were observed to be more disperse and closer to the tissue margins than correct classifications. The use of the RGB false color scheme provides more detail about the confidence of a specific classification. Classification of the 111,265 optical biopsies took a total of 23.56 seconds, or approximately 0.2 milliseconds per biopsy.

## 5 DISCUSSION

The study presented here points the way for a feasible approach to adding spectroscopic and depth-based scattering information derived from OCT scans to provide visual discrimination between selected types of tissue pathology in the colonic epithelium. The rationale for feasibility for clinical applciation is that with adequate training, the developed neural network was able to distinguish these tissue types quickly and with high accuracy. Our previous work with a mouse model of CRC [17] proposed the use of an optical biopsy consisting of binned OCT A-Scans as a means to better match the spatial resolution seen for traditional biopsies. This approach was further applied here by using 3x3 grids of A-scans to evaluate tissue properites. This provided better accuracy as it avoided the influence of an aberrant individual A-scan producing an incorrect classification. Instead, by examining larger localized regions, an accuracy of >93% was realized for distinguishing among the three tissue types studied here. However, if the goal is to simply identify histologically abnormal tissues, then upon grouping the tubular and tubulovillous tissue types, the accuracy achieved is near 98%.

The use of optical property information and neural network classification to provide visual discrimination can be an important factor for clinical translation. There is a premium placed on efficiency of colonoscopy procedures [24, 25] such that addition of novel microscopy methods [26] although potentially fertile sources of information, are often not widely adopted [27] because of their impact on procedure time. Here we have shown several schemes for creating visual contrast using optical properties. First, the attenuation coefficient has already been shown to provide contrast [15]. However, here we also note that derived parameters such as standard deviation and goodness of fit can also be used as a basis for contrast. Upon developing our neural network classification scheme, this information was also applied as a means to generate contrast. This approach showed that tissue type can rapidly be ascribed to localized regions of tissues imaged with OCT. The data are spectroscopically analyzed and fed into the developed neural network. The predicted tissue type is then indicated using a false color scheme to provide visual feedback. The speed of the neural network classifier suggests it could be used to provide this feedback in real time with further development.

Although this study shows promise, a significant limitation is the number and type of tissues examined. Initially, the protocol only sought to examine tissues extracted via physical biopsy which led to a preponderance of abnormal tissue samples. In later sessions, a priority was placed on also acquiring normal mucosa tissue biopsies from the same patient but at a distance from the observed polyps. Unfortunately, not all participants agreed to this additional biopsy. Further, only a few unique polyp morphologies (tubulovillous and tubular microstructural morphologies) were analyzed for this study. Although a handful of tissues of other types were observed, including sessile and laterally spreading bulk polyp morphologies, these were only represented by a single sample and the images provided a poor basis for training the algorithms. A broader imaging cohort could overcome these factors and also provide a means to disentangle information unique to tissue health from information unique to the patients themselves. Additionally, examination of a wider range of tissue pathologies could better illuminate the relationship between optical features and malignant potential. In a broader study, additional information such as polyp size, location in the colon, and patient demographics could also be examined to understand their influence on optical properties.

The approach developed here began with examination of individual A-scans and then progressed to analysis for 3x3 regions as an optical biopsy. This is meant to lessen the impact of individual A-scans which may be influenced by noise such as hyperreflection or signal dropout. However, this approach can also enable meaningful results to be obtained if a given tissue sample contains a range of tissue morphologies. In histopathology analysis of colon tissue biopsies, a sample is typically only given a single diagnosis. Further, the differences between classifications can be subjective. Without further detailed analysis to included consensus diagnoses and better define tissue margins, it is uncertain if this would be the correct classification for every portion of a volumetric image. The images in Figure 10 suggest that there may be a heterogeneity that must be considered. The false color images of the normal mucosa tissue samples show that there is potential for misclassification of each optical biopsy point. However, the classification is predominantly correct with 79%-91% of the pixels matching the normal mucosa histology diagnosis. While there is a chance that there may be some irregular tissue in the remaining 9%-21%, instead it seems that these misclassifications point to the achievable accuracy of the method as limited by variations in signal. In contrast the false color images of the tubular tissue samples show a bit lower accuracy of 61%-80%, with the remaining 20%-39% identified as normal mucosa. This suggests that a threshold of >50% of a given tissue type could be used to guide biopsy site selection.

The accuracy of spectroscopic OCT in classifying colon tissue shows promise for clinical applications. The high negative predictive value of 96.3% in the risk-based dichotomy indicates potentially utility in that doctors may not need to remove a polyp if they are able to determine in vivo that the sample is low risk, with potential benefits of reduced procedure time, procedure risk of performation, and reduced associated cost of tissue processing and analysis. If the current colonoscopy workflow does not allow for *in situ* analysis, tissues of interest can be excised and analyzed with spectroscopic OCT *ex vivo* and still produce the advantages of a quantitative classification scheme that does not require time-consuming and costly tissue staining, processing and histopathology analysis procedures. The high sensitivity and specificity in separating abnormal from normal mucosa tissue may provide utility in identifying regions of concern and even lead to early detection by observing features that are not seen visual inspection alone. Although on its own spectroscopic OCT may not yield sufficiently high sensitivity or positive predictive value in the risk-based dichotomy for clinical practice to rely on this metric by itself, our study suggests it could be a useful tool when used in conjunction with white light visual inspection of polyps or the margins of suspicious lesions.

Future work will enable rapid classification of colon tissue towards replacing the traditional standard of care processing and pathological analysis of tissues to diagnose disease, whether *ex vivo* after a suspicious sample has been resected, or in real time in an endoscopic probe. Additional potential applications of this work include eliminating the unnecessary resection of normal mucosa tissue or hyperplastic polyps, with benefits in reduction of procedure time and lower risk of complications due to resection, and margin assessment to get a better understanding of the efficacy of resection.

## 6 CONCLUSION

In conclusion, the feasibility of using optical properties derived from SOCT images has been evaluated using *ex vivo* colonic epithelium tissue. Parameters related to the attenuation of signal with depth and its spectral dependence were analyzed using LSTM and KNN machine learning approaches. While these produced some discrimination, a higher accuracy was obtained by using this information as input to a neural network classifier. Use of this information to generate contrast using false coloring shows a potential use of this approach for clinical guidance for identifying abnormal tissues in the colonic epithelium The potential clinical utility of this SOCT approach may be during colonoscopy procedures for CRC screening where real time definition of adenomatous tissue would provide the opportunity for immediate diagnosis. This could improve therapeutic procedures while also potentially reducing expense and time cost associated with further procedures. However, adoption for clinical use would require larger studies that characterized the optical properties of a wider range of adenoma and normal mucosa tissue morphologies.

## Abbreviations

CRC: colorectal cancer
KNN: K-nearest neighbor
LSTM: long short-term memory
OCT: optical coherence tomography
SOCT: spectroscopic optical coherence tomography
STFT: short time Fourier transform
SP: scattering power
FWHM: full width half maximum
CCD: charge-coupled device
ROC: receiving operator characteristic

## ACKNOWLEDGMENTS

NSF GRFP (WK), NIH STTR 1R41EB032693

## CONFLICT OF INTEREST

A.W. is President of Lumedica Inc.

## DATA AVAILABILITY STATEMENT

The data that support the findings of this study are available from the corresponding author upon reasonable request.

## Notes

### Competing Interest Statement

Adam Wax is founder and president of Lumedica, Inc. and Lumedica Vision

